# Description of *Klebsiella indica* sp. nov., isolated from the surface of tomato

**DOI:** 10.1101/708925

**Authors:** Sukriti Gujarati, Diptaraj Chaudhari, Mitesh Khairnar, Yogesh Shouche, Praveen Rahi

**Author notes:** **Corresponding author details:** Praveen Rahi, National Centre for Microbial Resource, National Centre for Cell Science, Pune, Maharashtra 411007, India, Email ID. The draft genome sequence has been deposited in GenBank under the accession number VCHQ00000000. The strain TOUT 106 is deposited with the culture collection at the National Centre for Microbial Resource, under the accession number MCC 2901.

## Abstract

A novel bacterial strain designated TOUT106^T^ was isolated from the surface of a tomato collected from the local vegetable market in Pune, India. The cells were rod shaped, Gram-stain-negative, encapsulated and non-motile. The strain TOUT106^T^ grows as mucoid and translucent colonies on blood agar medium and the best growth was observed at 28°C and at pH 7.0, and could tolerate up to 2% (w/v) NaCl. On the basis of 16S rRNA gene sequence analysis, strain TOUT106^T^ was placed under *Salmonella* clade, with close similarity to *Salmonella enterica* subsp. *arizonae* strain NCTC 8297^T^ (98.42%). Genome-based phylogenetic analysis revealed that the strain forms a distinct branch within the *Klebsiella* clade and *K. michiganensis* DSM25444^T^ and *K. oxytoca* NBRC105695^T^ were the closest neighbor. The genomic DNA G+C content of strain TOUT106^T^ was 53.53 mol%. The average nucleotide identity of TOUT106^T^ was less 86.4% with closely related members of the family *Enterobacteriaceae*. The major fatty acids of strain TOUT106^T^ were C_16:0_, C_17:0_ cyclo, C_14:0_ 3OH/C_16:1_ iso, C_14:0_, C_19:0 cyclo w8c_, C_18:1_ w6c/C_18:1_ w7c, C_12:0_ and C_16:1_ w7c/C_16:1_ w6c. The strain TOUT106^T^ showed differences in physiological, phenotypic and protein profiles by MALDI-TOF MS to its closest relatives. Based on the phenotypic including chemotaxonomic properties and phylogenetic analysis the strain TOUT106^T^ could be distinguished from the recognized species of the genus *Klebsiella*, was suggested to represent a novel species of this genus, for which the name *Klebsiella indica* sp. nov. is proposed. The type strain is TOUT106^T^ (=MCC 2901^T^).

## Introduction

The genus *Klebsiella* as defined by Trevisan in 1885 [1] with the type species *K. pneumoniae* [2] is polyphyletic [3] consisting of over 23 species of rod-shaped, Gram-stain-negative and capsule-forming bacteria. Recently, four novel species of the genus have been described viz., *K. michiganensis* [4], *K. quasipneumoniae* [5], *K. africanensis* [6] and *K. huaxiensis* [7] and the strains previously identified as *Klebsiella oxytoca* phylogroup Ko6 have been reclassified into a new species *K. grimontii* [8]. Members of this genus have been isolated from varied sources including clinical specimens [5, 7–9], soil [10, 11], forest swamps [12], water sample [13], household toothbrush holder [4] and mammalian gut [14]. *Klebsiella* isolates have also been reported frequently from various fruits and vegetables like sweet potato [15], banana [16], tomato [17], lettuce and cucumber [18]. While identifying a large collection of bacteria isolated from washed and unwashed fresh vegetables by using MALDI-TOF MS, we could not identify strain TOUT106, further analysis based on 16S rRNA gene sequencing revealed that the strain represents a putative new species. The aim of this study is to provide a detailed taxonomic description of a novel strain, TOUT106^T^, of the genus *Klebsiella* isolated from the washing of tomato surface.

## Material and methods

Strain TOUT106^T^ was isolated from washing the outer surface of tomato collected from a local vegetable market in Pune, India. The strain was inoculated on tryptone soya agar (TSA, Himedia M290) plates that were incubated at 28°C. The colonies were subcultured on TSA before preserving in 20 % (v/v) glycerol suspension at −80°C. The pure culture was processed for MALDI-TOF MS based identification, and the comparison of MALDI-TOF MS spectrum of TOUT106^T^ with the Biotyper 3.0 database resulted in no reliable identification. High-quality genomic DNA was extracted from the strain following the JGI protocol version 3 for bacterial genomic DNA isolation using CTAB [19]. The 16S rRNA gene sequence was amplified using universal primers (27f: 5’-AGAGTTTGATCCTGGCTCAG-3’ and 1492r: 5’-TACGGCTACCTTGTTACGACTT-3’) according to the methods described by Gulati *et al*. [20] and the amplified product was directly sequenced using the ABI PRISM Big Dye Terminator v3.1 Cycle Sequencing kit on a 3730xl Genetic Analyzer (Applied BioSystems). The similarity search for the 16S rRNA gene sequence of strain TOUT106^T^ was performed against the type strains of prokaryotic species in the EzBioCloud’s database [21]. Multiple alignment of sequences of strain TOUT106^T^ and its nearest neighbors were performed using the clustalW program in MEGA software v7.0 [22]. The aligned sequences were used to construct a phylogenetic tree with the Neighbor-joining algorithm and the topologies of the phylogenetic tree were assessed by bootstrap values based on 1000 replications. The 16S rRNA gene sequence of *Stenotrophomonas maltophilia* IAM 12423^T^ was used as outgroup.

Genome sequencing was performed on the Illumina MiSeq platform with 2 × 250 bp v2 chemistry and the good paired reads were subjected to de-novo assembly using MIRA version V5rc1 [23]. 16S rRNA gene sequence was retrived using the tool RNAmmer v1.2 [24]. The genome-based phylogenetic tree was reconstructed by both the FastTree [25] and RAxML [26] algorithm using the PATRIC v3.5.38 web service (https://www.patricbrc.org). The genomic data for all reference strains was downloaded from PATRIC database [27]. The Average Nucleotide Identity (ANI) was determined using the alignment-free sequence mapping approach between strain TOUT106^T^ and closely related strains of the *Enterobacteriaceae* family using FastANI [28] and a heatmap representation for the calculated ANI values was constructed using DisplayR (https://www.displayr.com/). Further pairwise genome-to-genome comparison in silico was made by calculating digital DNA-DNA hybridization (dDDH) by the recommended formula 2 using GGDC v2.1 [29]. Whole genome sequences were annotated using RAST [30] web server (http://rast.nmpdr.org/rast.cgi) and PATRIC v3.5.38 web service (https://www.patricbrc.org). The resistance gene identifier (RGI) v5.0.0 program was used to predict the presence of antimicrobial resistance genes using the comprehensive antibiotic resistance database (CARD) v3.0.2 and the PATRIC genome annotation service [31, 32]. The virulence factors in the strain TOUT106^T^ were predicted using the annotation data obtained by RAST as well as the Virulence Factor Database (VFDB) of pathogenic Bacteria using the VFAnalyzer tool [33].

For analysis of chemotaxonomic features, the strain was grown on TSA at 28 °C and cell biomass was harvested after 24 h unless otherwise stated. Preparation and analysis of fatty acid methyl esters were performed as described by Sasser [34] using the Microbial Identification System (MIDI) and the Microbial Identification software package (Sherlock version 6.1; MIDI database, TSBA6). To generate the Mean Spectral Profile (MSP) for the whole-cell protein ranging from 2-20 KDa, proteins were extracted using ethanol/formic acid after 24 h growth on TSA and the extract was analyzed by matrix-assisted laser desorption/ionization time-of-flight mass spectrometer (MALDI-TOF MS) autoflex speed (Bruker Daltonik GmbH) [35]. The newly generated MSP of strain was compared with those of the reference strains of closely related species present in the Bruker Biotyper database and principal component analysis (PCA) dendrogram was generated by using Biotyper 3.1.

Morphological, physiological and biochemical test for strain TOUT106^T^ and colony morphology was observed on TSA plates incubated under aerobic conditions. The strain was also incubated under anaerobic conditions in a Forma Anaerobic System Glovebox (Thermo Scientific, 1025). The growth pattern of the isolate was also determined on *Salmonella-Shigella* Agar plate (Himedia, M108D), MacConkey’s Agar plate (Himedia, M083) and sheep blood agar medium (Himedia, MP1301) supplemented with freshly procured sheep blood. Scanning electron microscopy was performed to observe cell morphology as described in Rahi *et al*. [20]. The cell morphology was observed on a scanning microscope (S3400N, Hitachi) at an acceleration voltage of 30.0 kV to determine the cell morphology and size. Gram-staining and capsule-staining kits (Himedia, K001 and K004) were used following manufacturer’s instructions. Cell motility was confirmed by hanging drop technique. Oxidase disc (Himedia, DD018) was used for testing oxidase activity and catalase activity was determined by bubble formation in a 3□% (v/v) H_2_O_2_ solution. Growth at different temperatures (4, 10, 15, 20, 28, 37, 45 and 55 °C), NaCl concentrations [0-5% (w/v) at 0.5% intervals] and pH values (4.0-11.0 at 1.0 pH unit intervals) was examined after incubation in tryptone soya broth (TSB) medium for 7 days in automated microbial growth analyzer (Bioscreen C, OY Growth Curves, Finland). The initial pH of the inoculation broth was adjusted using 1 M HCl and 1 M NaOH. Biochemical characteristics, enzyme activities, and oxidation/or reduction of carbon sources were performed using the API 20E and API ZYM systems (bioMérieux, 07584D and 25200) and Biolog GN III system following manufacturers’ instructions.

Tolerance to antibiotics was determined using the disc diffusion method (HiMedia, DE007, and DE001) after swabbing the cell suspension of turbidity equivalent to 0.5 McFarland standard on Mueller Hinton Agar (HiMedia, M1084) plates, and inhibition zone diameter was measured after 24 hours of incubation at 28°C [36–38]. The minimum inhibitory concentration (MIC) of the strain TOUT106^T^ was determined for colistin, ampicillin, cefotaxime, and ciprofloxacin in duplicates using the broth dilution method. Stock solutions were prepared as per the method described by J.M. Andrews [39]. Two-fold dilutions of test antibiotics were prepared in the range of 128-0.25 μg ml^-1^ for colistin and 256-0.5 μg/ml for ampicillin, cefotaxime and ciprofloxacin in sterilized Mueller-Hinton broth in sterile, flat-bottomed 96-well microtitre plates with lid (Falcon, 353072). The plates were inoculated with freshly grown bacterial culture of turbidity equivalent to 0.5 McFarland standard. The inoculated medium without the antibiotics served as the positive control while the uninoculated sterilized medium with the antibiotic served as the negative control. The plate was incubated at 37°C for 20 h after which 5 μl of resazurin indicator solution (5 mg/ml) was added to each well and the plate was further incubated for 2 h. The lowest antibiotic concentration, which prevented color change from purple to pink was recorded as MIC. The susceptibility to antibiotics was interpreted based on the Clinical and Laboratory Standards Institute (CLSI) guidelines determined for members of the family *Enterobacteriaceae* [36].

## Results

The sequence of 1392 bp was acquired by amplification and sequencing of 16S rRNA gene for the strain TOUT106^T^. However, two sequences of 16S rRNA gene were retrieved from the assembled draft genome (1528 and 1510 bp respectively) showing 97.16% similarity (98% query coverage) with each other, while 100% (100% query coverage) and 97.70% (100% query coverage) similarity with the acquired 1392 bp gene sequenced by Sanger’s method. Phylogenetic tree constructed on the basis of 16S rRNA gene sequences (1392 bp, 1510 bp and 1528 bp), placed the strain TOUT106^T^ within the *Salmonella* clade (Fig. S1), with close similarity to *Salmonella enterica* subsp. *arizonae* strain NCTC 8297^T^ (98.42 %).

The genome of TOUT106^T^ had a size of 5.24 Mbp with 86 final assembled contigs. The overall genome sequencing coverage of 164.25x and an N50 value of 191451 bp was obtained. A distinct branch was obtained for strain TOUT106^T^ within the *Klebsiella* clade and *K. michiganensis* DSM25444^T^ and *K. oxytoca* NBRC105695^T^ showed the closest genome-based phylogenetic affinity with the strain TOUT106^T^ (Fig 1). The genomic DNA G+C content of strain TOUT106^T^ was 53.53 mol%, which is well within the range (i.e. 53-58 mol %) of the genus *Klebsiella* [1]. The ANI value for the strain TOUT106^T^ as compared with other related members of the family *Enterobacteriaceae* was <86.4% (Fig. 2), while the dDDH relatedness of these strain was <70%, suggesting the strain TOUT106^T^ is novel species [27, 28, 40–42]. A detailed overview with related type strain genomes of the *Enterobacteriaceae* family used for comparison is given in Table 1.

**Fig. 1.**
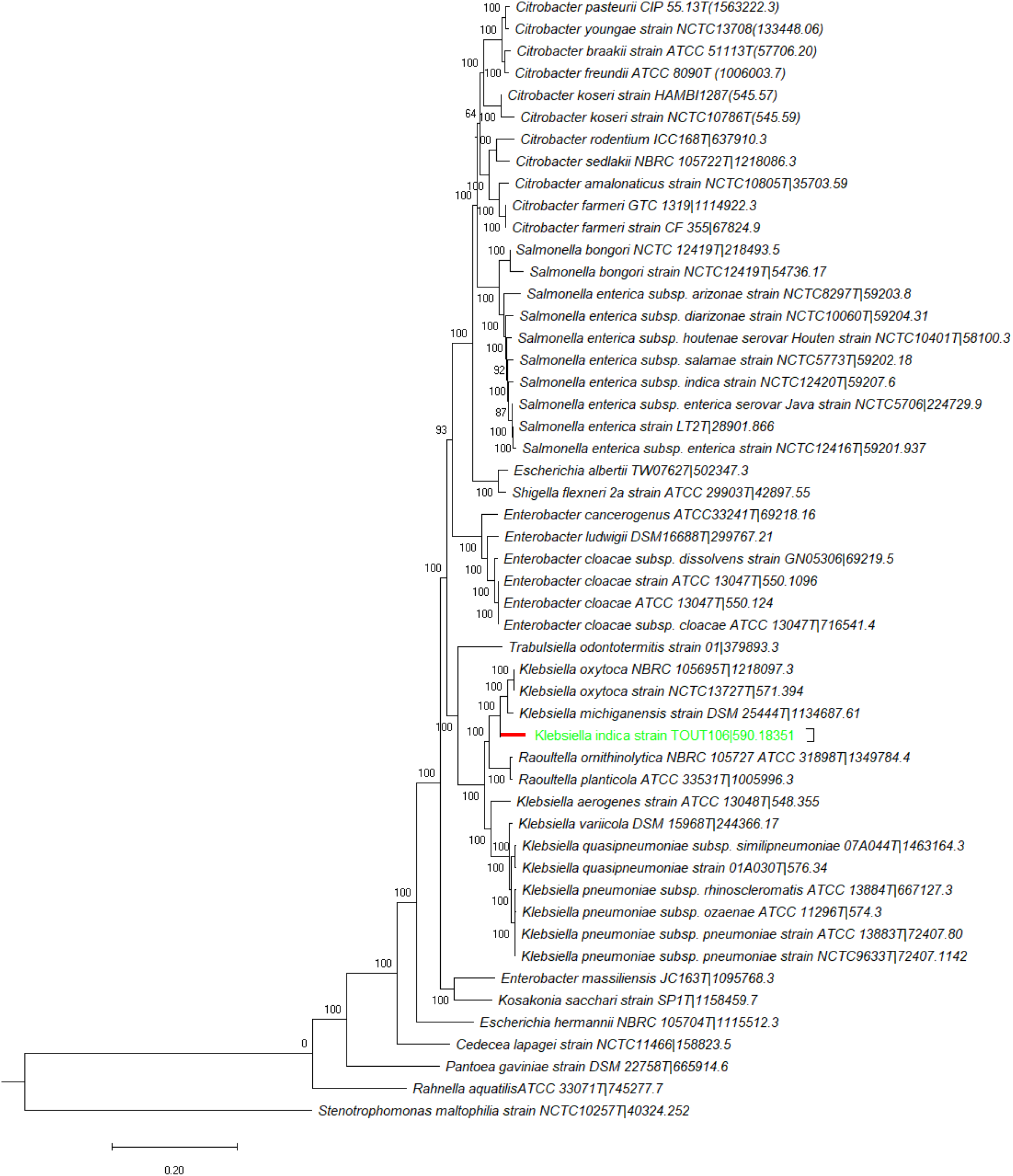
Genome based phylogenetic tree of the strain TOUT106^T^ (branch highlighted in red) with related strains of the family *Enterobacteriaceae* using the FastTree algorithm. *Stenotrophomonas maltophilia* NCTC10257^T^ was used as outgroup.

**Fig. 2.**
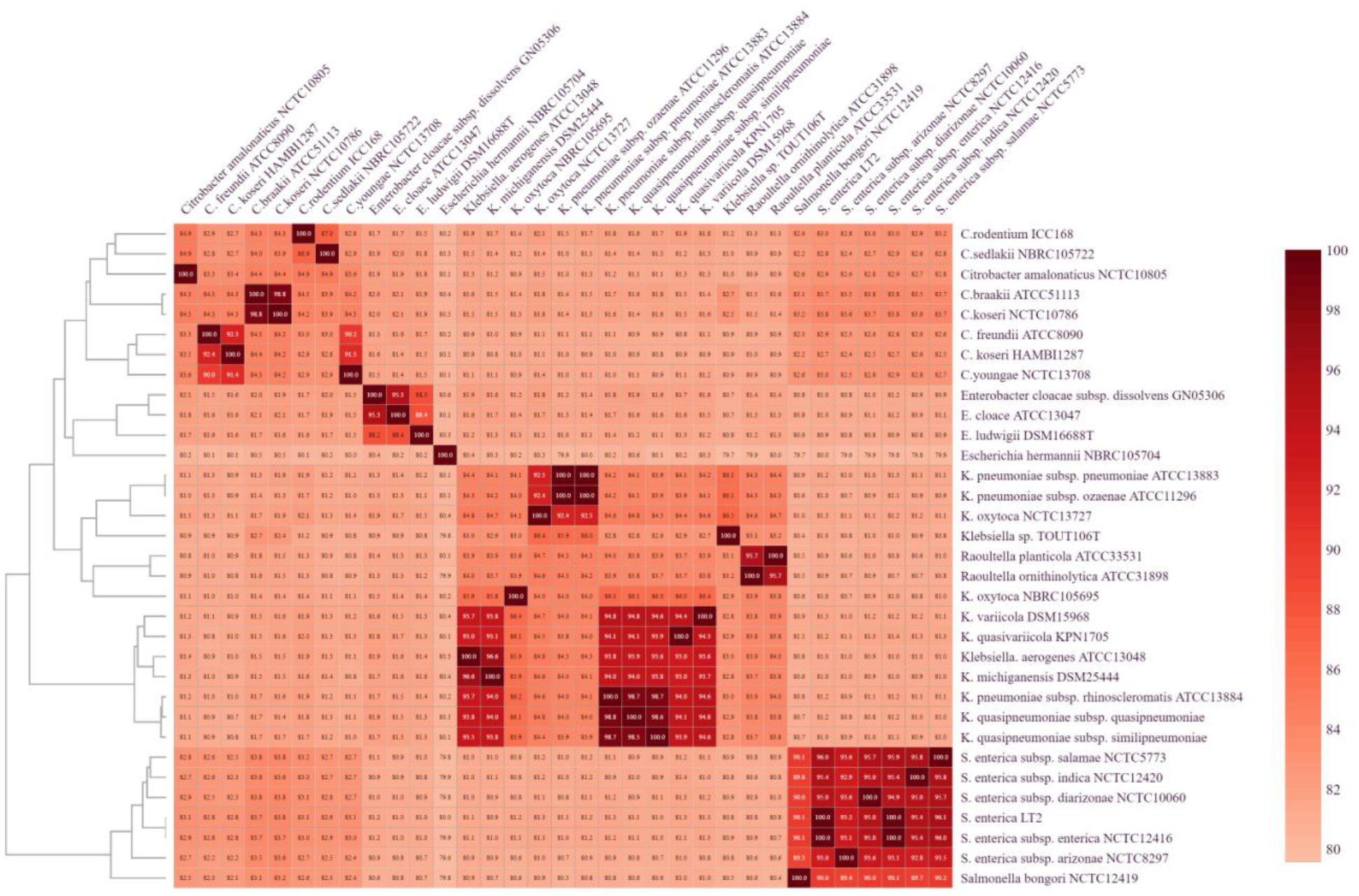
Heatmap and dendrogram of ANI values of the strain TOUT106^T^ with related type strains of the family *Enterobacteriaceae*.

**Table 1:**
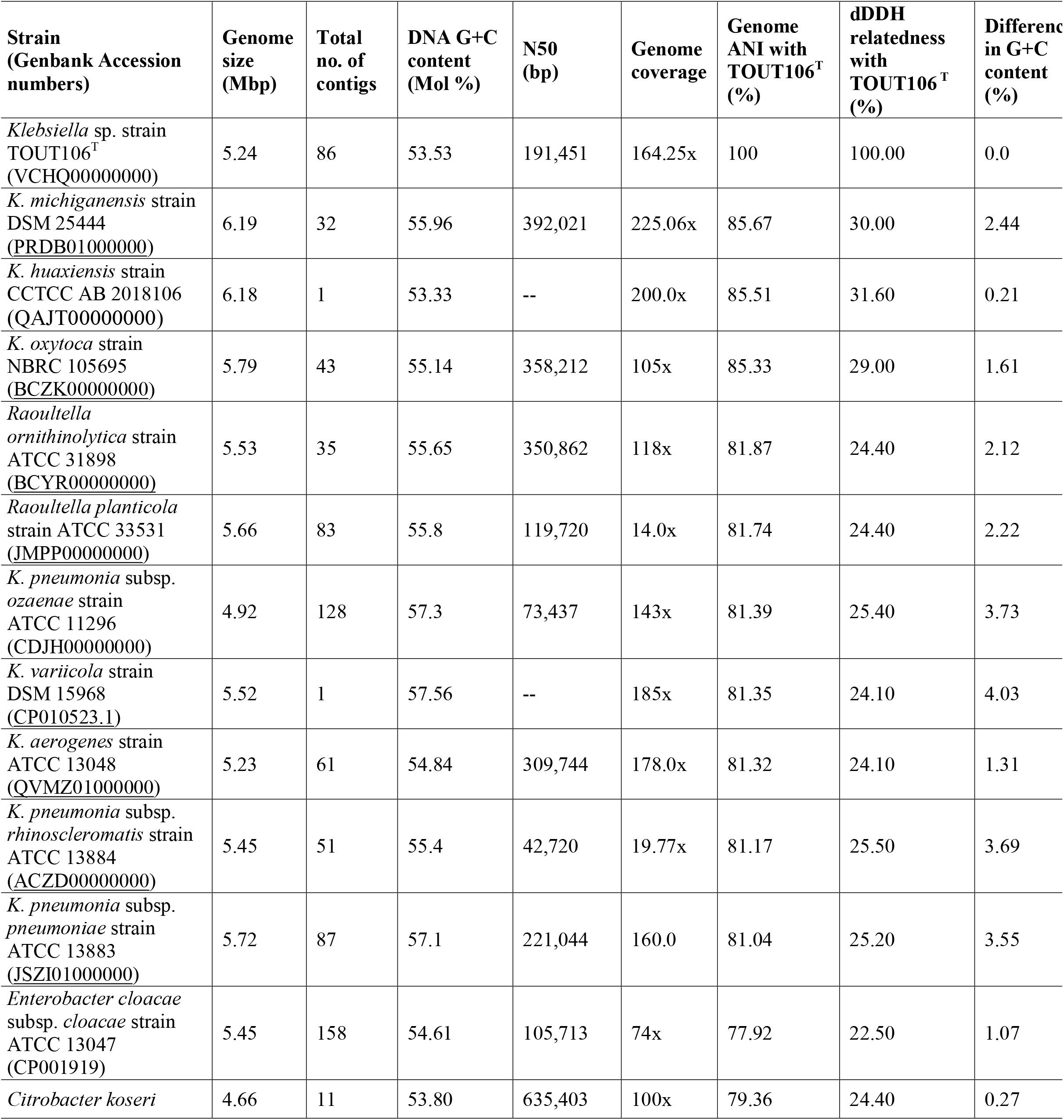
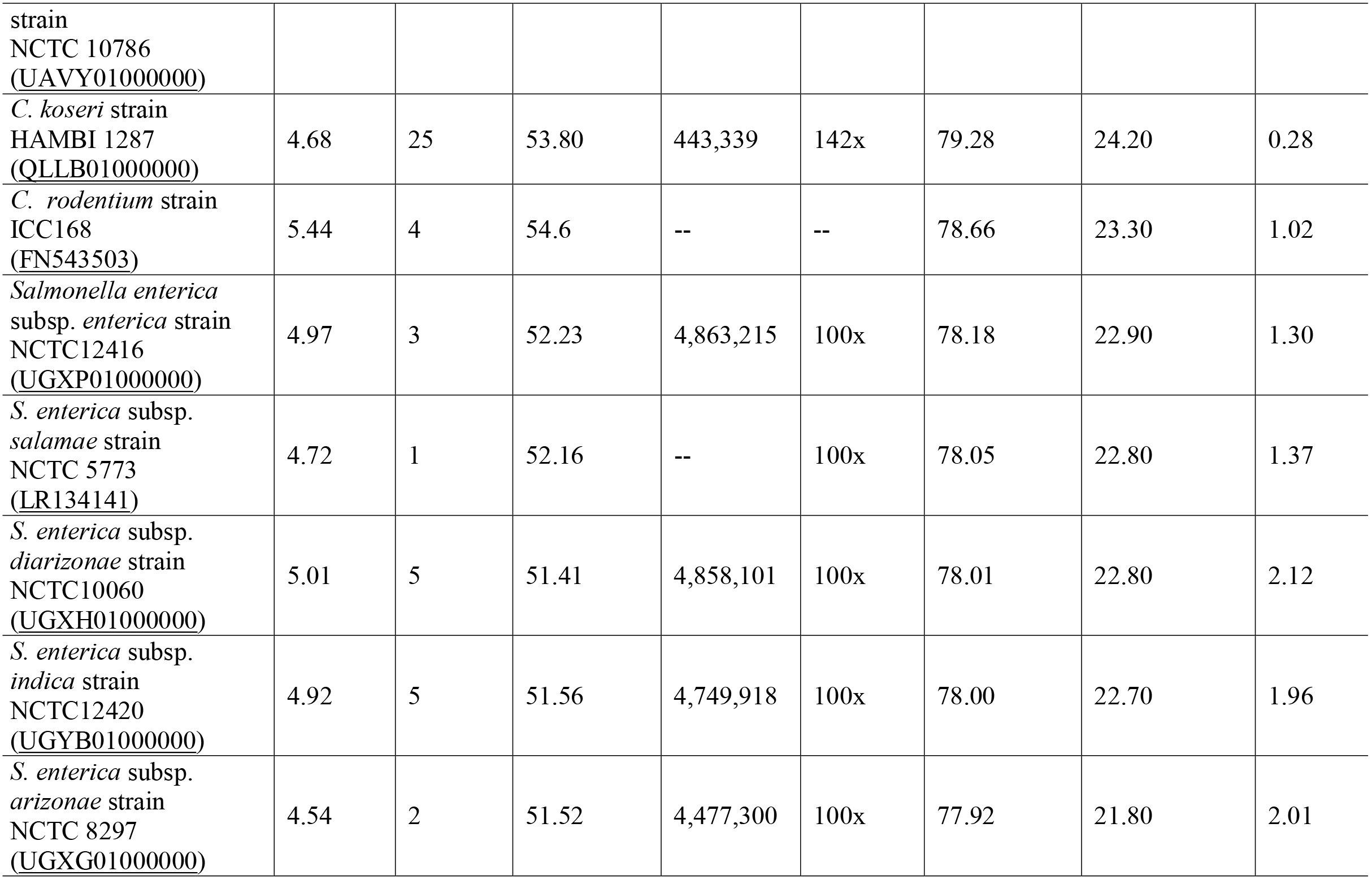
Genomic characterization of strain TOUT106^T^ with related type strains of the *Enterobacteriaceae* family.

| Strain<br>(Genbank Accession<br>numbers) | Genome<br>size<br>(Mbp) | Total<br>no. of<br>contigs | DNA G+C<br>content<br>(Mol %) | N50<br>(bp) | Genome<br>coverage | Genome<br>ANI with<br>TOUT106<sup>T</sup><br>(%) | dDDH<br>relatedness<br>with<br>TOUT106<sup>T</sup><br>(%) | Differenc<br>in G+C<br>content<br>(%) |
| --- | --- | --- | --- | --- | --- | --- | --- | --- |
| *Klebsiella* sp. strain<br>TOUT106<sup>T</sup><br>(VCHQ000000000) | 5.24 | 86 | 53.53 | 191,451 | 164.25x | 100 | 100.00 | 0.0 |
| *K. michiganensis* strain<br>DSM 25444<br>(PRDB010000000) | 6.19 | 32 | 55.96 | 392,021 | 225.06x | 85.67 | 30.00 | 2.44 |
| *K. huaxiensis* strain<br>CCTCC AB 2018106<br>(QAJT000000000) | 6.18 | 1 | 53.33 | -- | 200.0x | 85.51 | 31.60 | 0.21 |
| *K. oxytoca* strain<br>NBRC 105695<br>(BCZK000000000) | 5.79 | 43 | 55.14 | 358,212 | 105x | 85.33 | 29.00 | 1.61 |
| *Raoultella*<br>*ornithinolytica* strain<br>ATCC 31898<br>(BCYR000000000) | 5.53 | 35 | 55.65 | 350,862 | 118x | 81.87 | 24.40 | 2.12 |
| *Raoultella planticola*<br>strain ATCC 33531<br>(JMPP000000000) | 5.66 | 83 | 55.8 | 119,720 | 14.0x | 81.74 | 24.40 | 2.22 |
| *K. pneumonia* subsp.<br>*ozaenae* strain<br>ATCC 11296<br>(CDJH000000000) | 4.92 | 128 | 57.3 | 73,437 | 143x | 81.39 | 25.40 | 3.73 |
| *K. variicola* strain<br>DSM 15968<br>(CP010523.1) | 5.52 | 1 | 57.56 | -- | 185x | 81.35 | 24.10 | 4.03 |
| *K. aerogenes* strain<br>ATCC 13048<br>(QVMZ010000000) | 5.23 | 61 | 54.84 | 309,744 | 178.0x | 81.32 | 24.10 | 1.31 |
| *K. pneumonia* subsp.<br>*rhinoscleromatis* strain<br>ATCC 13884<br>(ACZD000000000) | 5.45 | 51 | 55.4 | 42,720 | 19.77x | 81.17 | 25.50 | 3.69 |
| *K. pneumonia* subsp.<br>*pneumoniae* strain<br>ATCC 13883<br>(JSZI010000000) | 5.72 | 87 | 57.1 | 221,044 | 160.0 | 81.04 | 25.20 | 3.55 |
| *Enterobacter cloacae*<br>subsp. *cloacae* strain<br>ATCC 13047<br>(CP001919) | 5.45 | 158 | 54.61 | 105,713 | 74x | 77.92 | 22.50 | 1.07 |
| *Citrobacter koseri* | 4.66 | 11 | 53.80 | 635,403 | 100x | 79.36 | 24.40 | 0.27 |
| strain<br>NCTC 10786<br>(UAVY01000000) |  |  |  |  |  |  |  |  |
| *C. koseri* strain<br>HAMBI 1287<br>(QLLB01000000) | 4.68 | 25 | 53.80 | 443,339 | 142x | 79.28 | 24.20 | 0.28 |
| *C. rodentium* strain<br>ICC168<br>(FN543503) | 5.44 | 4 | 54.6 | -- | -- | 78.66 | 23.30 | 1.02 |
| *Salmonella enterica*<br>subsp. *enterica* strain<br>NCTC12416<br>(UGXP01000000) | 4.97 | 3 | 52.23 | 4,863,215 | 100x | 78.18 | 22.90 | 1.30 |
| *S. enterica* subsp.<br>*salamae* strain<br>NCTC 5773<br>(LR134141) | 4.72 | 1 | 52.16 | -- | 100x | 78.05 | 22.80 | 1.37 |
| *S. enterica* subsp.<br>*diarizonae* strain<br>NCTC10060<br>(UGXH01000000) | 5.01 | 5 | 51.41 | 4,858,101 | 100x | 78.01 | 22.80 | 2.12 |
| *S. enterica* subsp.<br>*indica* strain<br>NCTC12420<br>(UGYB01000000) | 4.92 | 5 | 51.56 | 4,749,918 | 100x | 78.00 | 22.70 | 1.96 |
| *S. enterica* subsp.<br>*arizonae* strain<br>NCTC 8297<br>(UGXG01000000) | 4.54 | 2 | 51.52 | 4,477,300 | 100x | 77.92 | 21.80 | 2.01 |

The genome annotation revealed that the strain TOUT106^T^ showed the presence of 19 out of 20 nif genes (*nifA, nifB, nifD, nifE, nifH, nifJ, nifK, nifL, nifM, nifN, nifQ, nifS, nifT nifU, nifV, nifW,nifX, nifY, nifZ*,) involved in nitrogen fixation and regulation. The ability of the strain TOUT106^T^ for nitrogen fixation was further confirmed by growing it on nitrogen-free Jensen’s agar medium plate (Himedia, M710). Annotation of antimicrobial genes revealed that the resistome of strain TOUT106^T^ consists of genes conferring resistance to β-lactams (subclass B3 beta-lactamases, metallobeta-lactamases, PhnP protein, C-class beta-lactamases and PBP3), fosfomycin resistance (FosA5), macrolide resistance (*mdfA*), aminoglycoside resistance (*baeR*), and sulphonamide resistance (*folP*). The major resistance mechanisms employed by strain TOUT106^T^ include antibiotic target alteration, antibiotic efflux, reduced permeability to antibiotics and antibiotic inactivation, as predicted by CARD. Genes responsible for capsule synthesis (*galF, manB)*, regulation of capsule synthesis (*rcsA, rcsB*), lipopolysaccharide (WzzE, *msbA*), siderophores, including enterobactin (*febBCDG* locus*, entA, entB, entS and ybt locus*), aerobactin (iron-chelator utilizor protein), yersiniabactin (*fyuA, ybtA, ybtP, ybtQ, ybtX*) and type I (*fimA, fimC, fimD, fimG, fimH*) fimbriae were observed in strain TOUT106, which are major virulence factors reported in *Klebsiella* [43].

The major fatty acid detected in strain TOUT106^T^ was C_16:0_ (44.07%) that is typical for members of the genus *Klebsiella*. Other detected fatty acids include C_17:0 cyclo_ (15.325%), summed feature 2 (C_14:0_ 3OH/C_16:1_ iso, 9.62%), C_14:0_ (8.99%), C_19:0 cyclo w8c_ (6.14%), summed feature 8(C_18:1_w6c/C_18:1_ w7c, 5.1%), C_12:0_ (3.095%), summed feature 3 (C_16:1_ w7c/C_16:1_ w6c, 3.85%), and summed feature 5 (C_18:0_ ante/C_18:2_ w6, 9c, 1.17%). Lower amounts of summed feature 3 (C_16:1_ w7c/C_16:1_ w6c, 3.85%) and summed feature 8 (C_18:1_ w6c/C_18:1_ w7c, 5.1%) were detected in the strain TOUT106^T^, on the contrary these fatty acids were relatively high in reference strains *Klebsiella michiganensis* DSM 25444^T^ and *Klebsiella oxytoca* ATCC 13182^T^(Table 2). A comparison of MALDI-TOF MS spectra based dendrogram showed that the strain TOUT106^T^ clearly separated from the type strains of *Salmonella*, and was placed along with *Klebsiella variicola* DSM 15968^T^ and *Roultella terrigena* DSM 2687^T^, corroborating well with the results of genome-based analysis (Fig. S2).

**Table 2.**
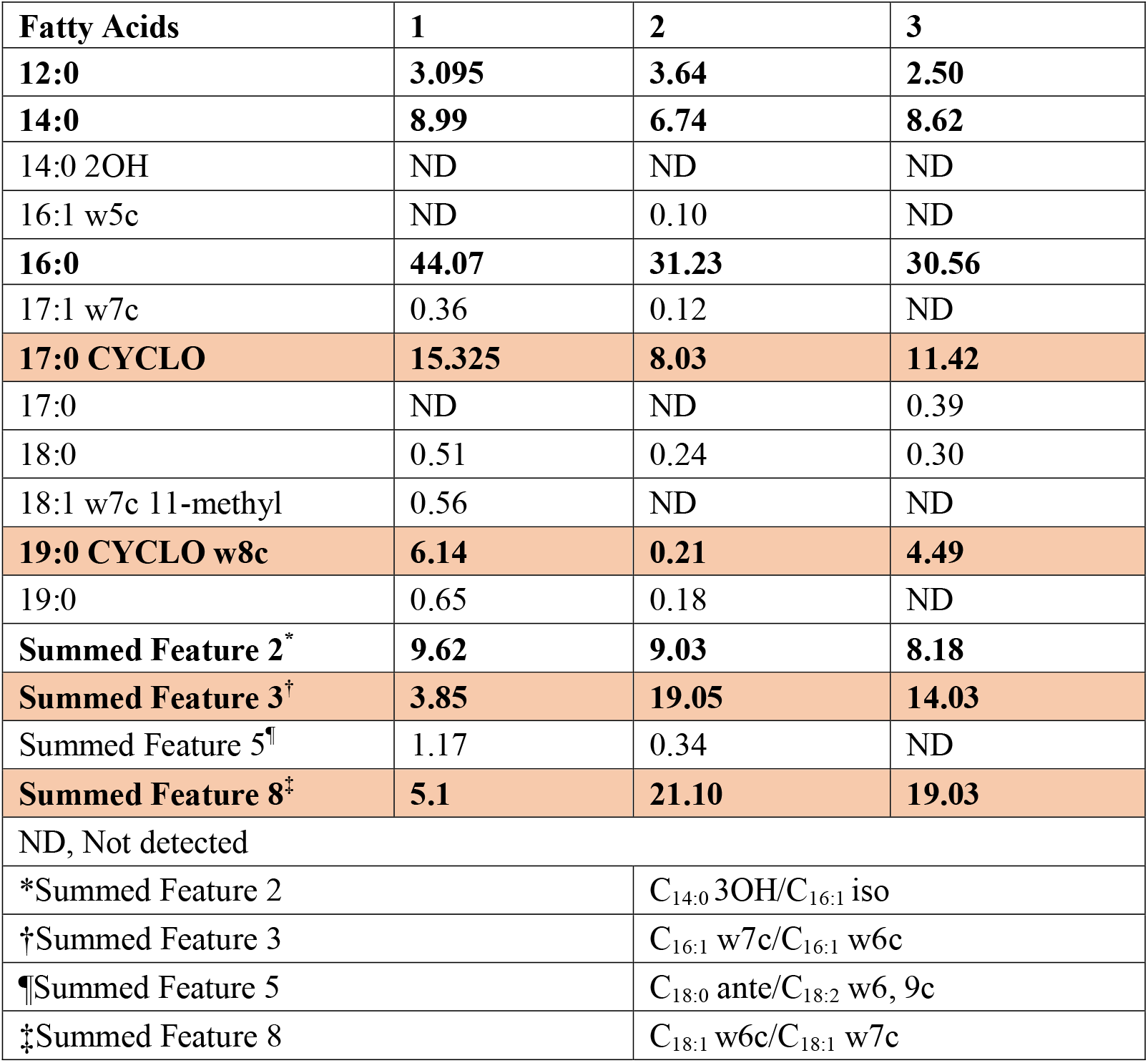
A comparative account of long chain fatty acid composition (%) of strain TOUT106^T^ with the closest phylogenetic neighbors [4]. Taxa: 1, TOUT106^T^, 2, *K. michiganensis* DSM 25444^T^, 3, *K. oxytoca* ATCC 13182^T^

| Fatty Acids | 1 | 2 | 3 |
| --- | --- | --- | --- |
| **12:0** | **3.095** | **3.64** | **2.50** |
| **14:0** | **8.99** | **6.74** | **8.62** |
| 14:0 2OH | ND | ND | ND |
| 16:1 w5c | ND | 0.10 | ND |
| **16:0** | **44.07** | **31.23** | **30.56** |
| 17:1 w7c | 0.36 | 0.12 | ND |
| **17:0 CYCLO** | **15.325** | **8.03** | **11.42** |
| 17:0 | ND | ND | 0.39 |
| 18:0 | 0.51 | 0.24 | 0.30 |
| 18:1 w7c 11-methyl | 0.56 | ND | ND |
| **19:0 CYCLO w8c** | **6.14** | **0.21** | **4.49** |
| 19:0 | 0.65 | 0.18 | ND |
| **Summed Feature 2<sup>\*</sup>** | **9.62** | **9.03** | **8.18** |
| **Summed Feature 3<sup>†</sup>** | **3.85** | **19.05** | **14.03** |
| Summed Feature 5<sup>¶</sup> | 1.17 | 0.34 | ND |
| **Summed Feature 8<sup>‡</sup>** | **5.1** | **21.10** | **19.03** |
| ND, Not detected |  |  |  |
| *Summed Feature 2 |  | C<sub>14:0</sub> 3OH/C<sub>16:1</sub> iso |  |
| †Summed Feature 3 |  | C<sub>16:1</sub> w7c/C<sub>16:1</sub> w6c |  |
| ¶Summed Feature 5 |  | C<sub>18:0</sub> ante/C<sub>18:2</sub> w6, 9c |  |
| ‡Summed Feature 8 |  | C<sub>18:1</sub> w6c/C<sub>18:1</sub> w7c |  |

Colony morphology, as examined on blood agar medium were mucoid and translucent. However, a negative string test indicated the strain to be non-hypermucoviscous [44] (Fig. S4). Morphological analysis revealed that strain TOUT106^T^ is a rod shaped, Gram stain negative, encapsulated, non-motile, oxidase negative and catalase weekly positive, can grow anaerobically bacteria with size ranging from 0.7-0.9×2-3 μm (Fig. S3). The strain TOUT106^T^ could grow at 20-37 °C (optimum 28°C), pH ranging from 4-10 (optimum 7.0) and NaCl concentration tolerance up to 2% (Table 3). Sole carbon source utilization test showed that out of 71 carbon substrates in the GENIII BIOLOG microplate, TOUT106^T^ could utilize 46 substrates, of which four were partially used (Table S2). The isolate can be distinguished from its closest phylogenetic neighbours on the basis of growth at 10 and 45°C, salt tolerance, urease activity, lysine decarboxylase activity, Voges-Proskauer and methyl red test (Table 3). Strain TOUT106^T^ failed to grow at 10 and 45°C while both the reference strains showed growth at 10 and 45°C. Strain TOUT106^T^ is negative for lysine decarboxylase activity and Voges-Proskauer test and positive for methyl red test, while the reference strains are positive for lysine decarboxylase activity and Voges-Proskauer test and negative for methyl red reaction. Strain TOUT106^T^ shows negative urease activity, while *K. michiganensis* DSM 25444^T^ is urease negative and *K. oxytoca* ATCC 13182^T^ is urease positive. The strain TOUT106^T^ was found to be resistant to cefpodoxime (10 mcg), augmentin (30 mcg) and amoxicillin (10 mcg) out of the antibiotics tested using the Kirby-Bauer disc diffusion method (Table S3). The MIC value of the strain TOUT106^T^ for colistin was 64 mcg/ml while it was susceptible to ampicillin, cefotaxime and ciprofloxacin (Table S4).

**Table 3.** Differential phenotypic characteristics of the strain TOUT106^T^ in comparison to closest phylogenetic neighbours Taxa: 1, TOUT106^T^, 2, *K. michiganensis* DSM 25444^T^ [4], 3, *K. oxytoca ATCC 13182^T^*[4][12].

| Characteristics | 1 | 2 | 3 |
| --- | --- | --- | --- |
| Isolation source | Outer surface of tomato | Household toothbrush holder | Human pharyngeal tonsil |
| Temperature range (°C) | 20-37 (optimum 28°C) | 10-45 (optimum 35°C) | ND |
| pH range for growth | 4.0-10.0 (optimum 7.0) | Upto 10.0 ( optimum 7.0) | ND |
| NaCl tolerance | 0.0-2.0 % | Upto 6% | ND |
| Growth at 10°C | - | + | + |
| Growth at 45°C | - | + | + |
| Urease production | - | - | + |
| Lysine decarboxylase | - | + | + |
| Voges–Proskauer | - | + | + |
| Methy red reaction | + | - | - |
| G+C content (mol %) | 53.53% | 54.6% | 55.14% |

ND: Not determined

Based on the genome-based phylogeny, fatty acids and DNA G+C content it is indicated that the strain TOUT106^T^ is a member of genus *Klebsiella*. However, it differs from closely related species of the genus *Klebsiella* in several aspects, such as 16S rRNA gene, biochemical features, physiological features, protein profile and overall genome relatedness indices. Thus, representing novel species of the genus *Klebsiella*, for which the name *Klebsiella indica* sp. nov. is proposed.

### Description of *Klebsiella indica* sp. nov

***Klebsiella indica*** (in·di·ca L. fem. adj. *indica*, of or belonging to India, where the type strain was isolated from outer wash of tomato).

Cells are Gram-stain-negative, straight rods with round ends (0.7-0.9×2-3 μm) and non-motile. Colonies grown on trypticase soy agar are 1-3 mm in diameter, circular, raised with an entire margin and translucent opacity. Optimal temperature for growth is 28 °C and optimal pH is 7.0. Growth occurs in the absence of NaCl with up to 2% tolerance in trypticase soy broth. Weekly positive for catalase and negative for oxidase activity. Strain showed positive results in Biolog GN III analyses for utilization of dextrin, d-maltose, d-trehalose, d-cellobiose, d-gentiobiose, sucrose, d-raffinose, a-d-lactose, d-melibiose, b-methyl-d-glucoside, d-salicin, n-acetyl-d-glucosamine, n-acetyl-d-galactosamine, a-d-glucose, d-mannose, d-fructose, d-galactose, pectin, L-fucose, L-rhamnose, inosine, sodium lactate, d-sorbitol, d-manitol, d-arabitol, myo-inositol, glycerol, d-glucose-6-PO4, d-fructose-6-PO4, d-serine, troleandomycin, rifamycin, glycyl-L-proline, L-aspartic acid, L-glutamic acid, l-serine, guanidine HCl, niaproof 4, d-galacturonic acid, L-galactonic acid lactone, d-gluconic acid, d-glucuronic acid, mucic acid, d-saccharic acid, vancomycin, L-lactic acid, citric acid, a-keto-glutaric acid, d,l-malic acid, methyl pyruvate, bromo-succinic acid, lithium chloride, sodium butyrate, tetrazolium violet and blue (Table S2). Positive results in API ZYM strips for alkaline phosphatase, leucine arylamidase, acid phosphatase, naphthol-AS-BI-phosphohydrolase, β-glucosidase activities (Table S5). From API 20E tests, positive results are obtained for ß-galactosidase activity, citrate utilization, indole production, glucose, mannitol, inositol, sorbitol, rhamnose, saccharose, melibiose, amygdain and arabinose fermentation/oxidation while negative results are obtained for arginine dihydrolyase, lysine decarboxylase, ornithine decarboxylase, urease tryptophan deaminase, gelatinase activities, H_2_S production and Voges-Prauskauer test. It is resistant to cefpodoxime, augmentin and amoxicillin but susceptible to cefotaxime, cephtriaxone, amikacin, ampicillin, streptomycin, gentamicin, ciprofloxacin, levofloxacin, norfloxacin, chloramphenicol, tetracycline, rifampicin, cefixime and cotrimazole. MIC value for colistin is reported to be 64 mcg/ml. C_16:0_ and C_17:0 cyclo_ are the predominant cellular fatty acids. The DNA G+C content of the type strain is 53.53 mol%. The type strain TOUT106^T^ (= MCC 2901^T^) was isolated from outer surface washing of tomato, in Pune, India. The GenBank sequence accession no. for genome sequence is VCHQ00000000.

## Acknowledgement

Authors acknowledge the Director, NCCS, Pune, and financial support by the Department of Biotechnology (BT/Coord. II/01/03/2016).

## Ethical Statement

The experiments reported in this manuscript did not involve human participants and/or animals.

