## Supplementary data for "Description of *Klebsiella indica* sp. nov., isolated from the surface of tomato"

**Running Title:** *Klebsiella indica* sp. nov.,

**Authors and affiliations:**

Sukriti Gujarati^1^, Diptaraj Chaudhari^1^, Mitesh Khairnar^1^, Yogesh Shouche^1^, Praveen Rahi^1^*

^1^National Centre for Microbial Resource, National Centre for Cell Science, Pune

**Corresponding author details**:

Praveen Rahi,

National Centre for Microbial Resource, National Centre for Cell Science, Pune, Maharashtra 411007, India

Table S1: Percent similarity between 16 S rRNA gene sequence and five other key housekeeping genes (*atpD, gyrB, infB, recA* and *rpoB*) of the test strain TOUT106^T^ with that of related strains of the family *Enterobacteriaceae*.

| **Strain** | **16S rRNA gene sequence** | | | **Key housekeeping genes**  **(retrieved from genome)** | | | | |
| --- | --- | --- | --- | --- | --- | --- | --- | --- |
| (Genbank accession number) | 1528 bp^*^ | 1510 bp^#^ | Amplified (1392 bp) | *rpoB* | *gyrB* | *infB* | *recA* | *atpD* |
| *Salmonella enterica* subsp. *enterica* strain NCTC12416 ([UGXP01000000](http://www.ncbi.nlm.nih.gov/nuccore/UGXP01000000)) | 97.26 | 98.28 | 97.87 | 92.70 | 89.46 | 88.69 | 88.56 | 94.50 |
| *Salmonella enterica* subsp. *diarizonae* strain NCTC10060 (UGXH01000000) | 97.65 | 98.61 | 98.07 | 93.05 | 89.90 | 87.64 | 88.49 | 93.42 |
| *Salmonella enterica* subsp. *indica* strain NCTC12420 ([UGYB01000000](http://www.ncbi.nlm.nih.gov/nuccore/UGYB01000000)) | 97.91 | 98.68 | 98.08 | 92.90 | 89.53 | 88.12 | 88.36 | 94.58 |
| *Salmonella enterica* subsp. *arizonae* strain NCTC 8297 ([UGXG01000000](http://www.ncbi.nlm.nih.gov/nuccore/UGXG01000000)) | 97.78 | 98.41 | 98.42 | 92.88 | 88.95 | 88.08 | 88.07 | 93.64 |
| *Salmonella enterica* subsp*. salamae* strain NCTC 5773 ([LR134141](http://www.ncbi.nlm.nih.gov/nuccore/LR134141)) | 97.32 | 98.21 | 97.91 | 92.65 | 90.28 | 88.08 | 88.07 | 94.22 |
| *Citrobacter rodentium* ICC168  ([FN543503](http://www.ncbi.nlm.nih.gov/nuccore/FN543502,FN543504,FN543505,FN543503)) | 96.80 | 97.88 | 97.12 | 93.00 | 90.49 | 88.14 | 88.47 | 93.93 |
| *Citrobacter koseri* NCTC 10786 ([UAVY01000000](http://www.ncbi.nlm.nih.gov/nuccore/UAVY01000000)) | 97.06 | 98.94 | 97.34 | 95.34 | 95.74 | 89.65 | 88.44 | 95.08 |
| *Enterobacter cloacae* subsp. cloacae ATCC 13047 (CP001919) | 97.78 | 97.36 | 97.85 | 92.33 | 88.87 | 87.69 | 88.98 | 93.64 |
| *Klebsiella aerogenes* strain ATCC 13048 ([QVMZ01000000](http://www.ncbi.nlm.nih.gov/nuccore/QVMZ01000000)) | 96.59 | 95.78 | 96.59 | 93.67 | 90.43 | 90.31 | 89.33 | 94.22 |
| *Klebsiella variicola* DSM 15968 ([CP010523.1](http://www.ncbi.nlm.nih.gov/nuccore/CP010523.1)) | 97.51 | 96.50 | 97.38 | 93.60 | 89.90 | 90.31 | 89.52 | 94.07 |
| *Klebsiella michiganensis* strain DSM 25444 ([PRDB01000000](http://www.ncbi.nlm.nih.gov/nuccore/PRDB01000000)) | 96.55 | 95.63 | 96.56 | 94.12 | 89.37 | 90.61 | 92.73 | 96.24 |
| *Klebsiella oxytoca* NBRC 105695 ([BCZK00000000](http://www.ncbi.nlm.nih.gov/nuccore/BCZK00000000)) | 97.75 | 97.37 | 97.43 | 93.99 | 88.99 | 91.01 | 92.92 | 96.31 |
| *Raoultella planticola* ATCC 33531 ([JMPP00000000](http://www.ncbi.nlm.nih.gov/nuccore/JMPP00000000)) | 96.40 | 95.90 | 96.32 | 93.60 | 89.20 | 90.25 | 89.74 | 94.36 |
| *Raoultella ornithinolytica* ATCC 31898 ([BCYR00000000](http://www.ncbi.nlm.nih.gov/nuccore/BCYR00000000)) | 96.09 | 95.55 | 96.34 | 93.72 | 89.25 | 89.99 | 89.30 | 94.36 |
| *Klebsiella pneumonia* subsp. *rhinoscleromatis* ATCC 13884 ([ACZD00000000](http://www.ncbi.nlm.nih.gov/nuccore/ACZD00000000)) | 96.66 | 95.83 | 96.98 | 93.37 | 89.99 | 90.42 | 89.05 | 94.14 |
| *Klebsiella pneumonia* subsp.*pneumoniae* ATCC 13883 ([JSZI01000000](http://www.ncbi.nlm.nih.gov/nuccore/JSZI01000000)) | 97.25 | 96.43 | 97.20 | 93.42 | 89.86 | 90.31 | 89.14 | 94.22 |
| *Klebsiella pneumonia* subsp. *ozaenae* ATCC 11296  [(CDJH00000000](http://www.ncbi.nlm.nih.gov/nuccore/CDJH00000000)) | 96.59 | 95.83 | 96.84 | 93.52 | 90.07 | 90.27 | 88.95 | 94.14 |

*Klebsiella oxytoca* JCM 1665T (AB004754)

*Klebsiella michiganensis* strain W14 (NR 118335.1)

*Klebsiella oxytoca* ATCC 13182 (NR 041749.1)

*Klebsiella aerogenes* KCTC 2190 (NR 102493.2)

*Raoultella planticola* ATCC 33531 (NR 119279.1)

R*aoultella ornithinolytica* strain ATCC 31898 (NR 114502.1)

*Citrobacter murliniae* CDC 2970-59T (AF025369)

*Citrobacter pasteurii*  CIP 55.13 (KP057683.1)

*Citrobacter braakii*  DSM 17596 (HG798904.1)

*Erwinia aphidicola* DSM 19347T (AB273744)

*Enterobacter ludwigii* DSM 16688T (AJ853891)

*Enterobacter cancerogenus* LMG 2693T (Z96078)

*Klebsiella pneumoniae* subsp. *ozaenae* ATCC 11296 (NR 041750.1)

*Klebsiella quasipneumoniae* subsp. *similipneumoniae* 07A044T (HG933295)

*K. quasipneumoniae* subsp. s*imilipneumoniae* CW-D 3 (NR 132596.1)

*Klebsiella pneumoniae* subsp. *rhinoscelromatis* ATCC13884T (Y17657.1)

*Klebsiella pneumoniae* subsp. *pneumoniae* JCM 1662 (AB594768.1)

*Enterobacter cloacae* subsp. *dissolvens* LMG 2683T (Z96079)

*Enterobacter* *cloacae* DSM 30054T (NR 117679)

*Enterobacter cloacae* subsp. *cloacae* ATCC 13047 (CP001918)

*Kosakonia sacchari* SP1T (JQ001784)

*Citrobacter youngae* strain GTC 1314 (NR 041527.1)

*Cedecea lapagei* GTC 346T (AB273742)

*Salmonella enterica* subsp. *enterica* LT2T (AE006468)

*Salmonella enterica* subsp. *salamae* DSM 9220T (EU014685)

*Salmonella enterica* subsp. *diarizonae* DSM 14847T (EU014688)

*Salmonella enterica* subsp. *indica* DSM 14848T (EU014680)

*Salmonella enterica* subsp. *arizonae* strain ATCC 13314T (NR041696)

*Klebsiella indica* TOUT106*

*Klebsiella indica* TOUT106^¶^  (VCHQ00000000)

*Citrobacter farmeri* strain CIP 104553 (KM515968.1)

*Citrobacter amalonaticus*  LMG 7873 (NR 118106.1)

*Citrobacter sedlakii* NBRC 105722 (AB682286.1)

*Trabulsiella odontotermitis* Eant 3-9T (DQ453129)

*Citrobacter koseri* strain CDC-8132-86 (NR 104890.1)

*Klebsiella indica* TOUT106^#^ (VCHQ00000000)

*Escherichia albertii* TW07627T (ABKX01000030)

*Escherichia coli* ATCC 11775T (X80725.1)

*Shigella flexneri* ATCC 29903T (X96963)

*Escherichia hermannii* GTC347T (AB273738)

*Pantoea gaviniae* A18/07T (GQ367483)

*Rahnella aquatica* strain DSM 4594 (AJ233426)

*Stenotrophomonas maltophilia* strain IAM 12423 (AB294553)

0.020

Fig. S1: Phylogenetic tree based on all the three 16S rRNA gene sequences (1392 bp, 1528 bp and 1510 bp) of the strain TOUT106^T^ constructed using the neighbor joining algorithm.

*: Query of 1392 bp amplified and sequenced using Sanger’s method; ^¶^: Query of 1528 bp retrieved from genome; ^#^: Query of 1510 bp retrieved from genome

Fig. S2: Dendrogram of strain TOUT106^T^ and closely related species based on their intact cell proteins (2-20 KDa) MALDI-TOF MS mean spectra.

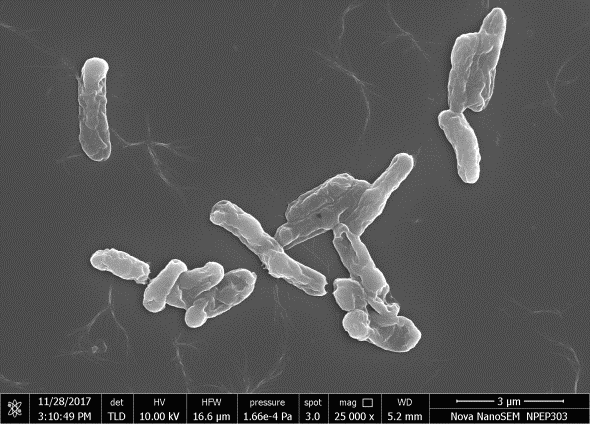

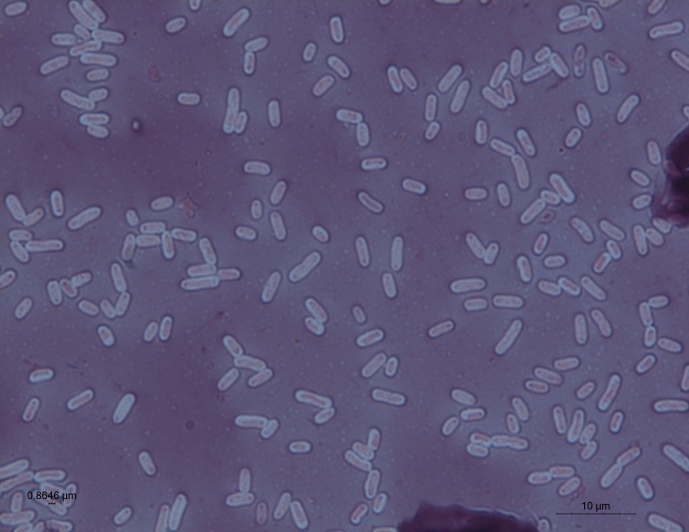

(b)

(a)

Fig. S3. Photomicrographs of strain TOUT106^T^, (a) scanning electron microscopy and (b) capsule staining.

(a)

(b)

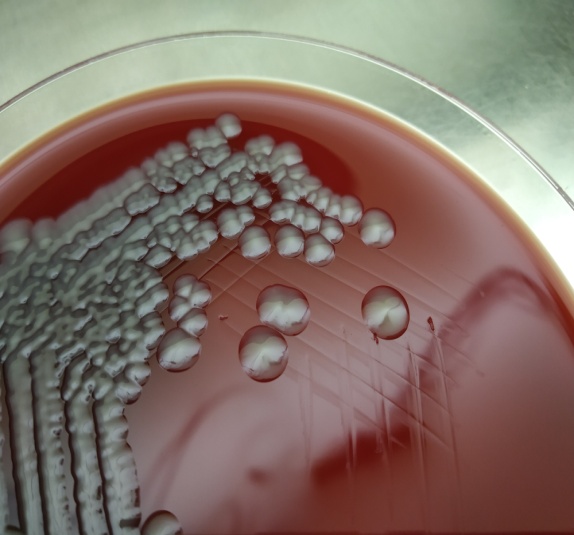

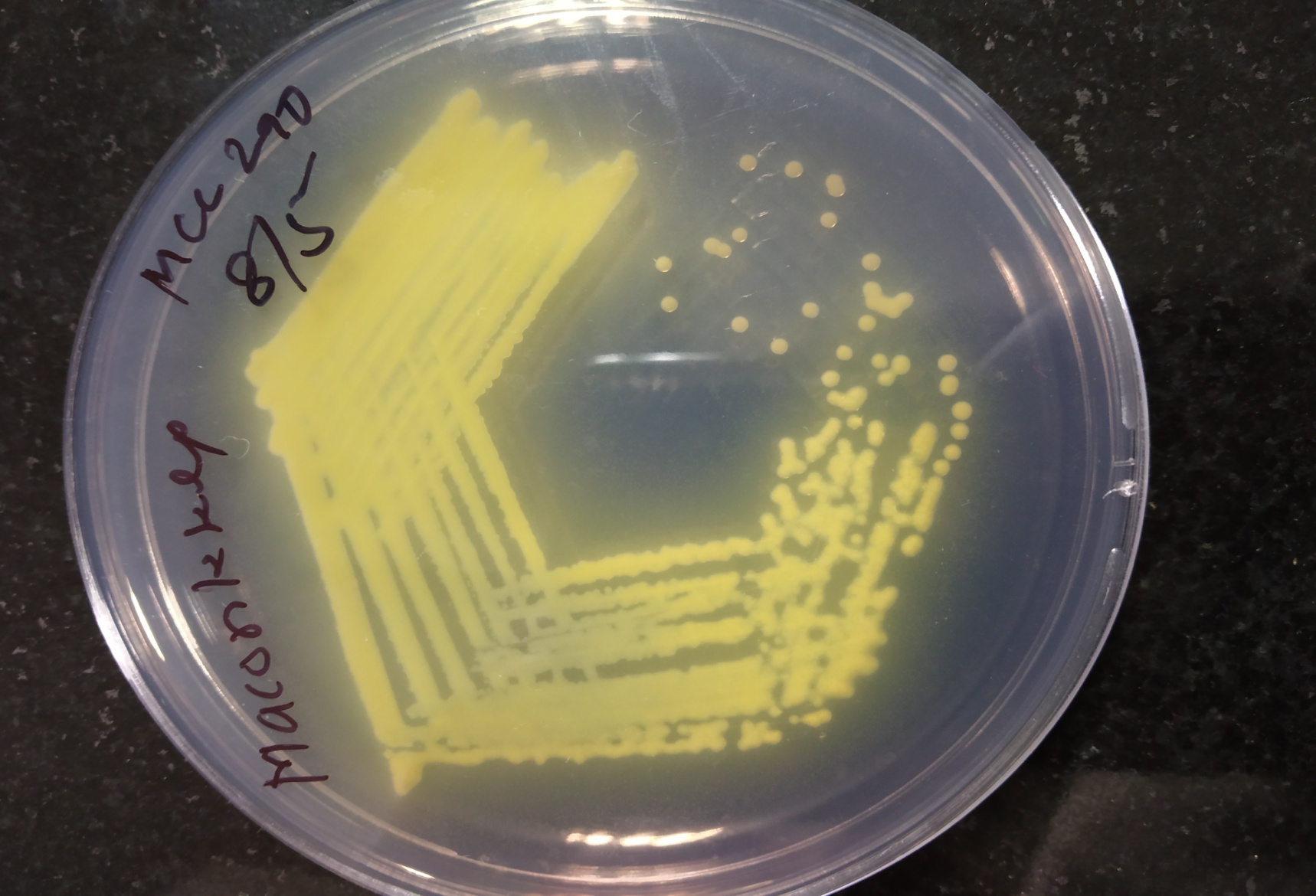

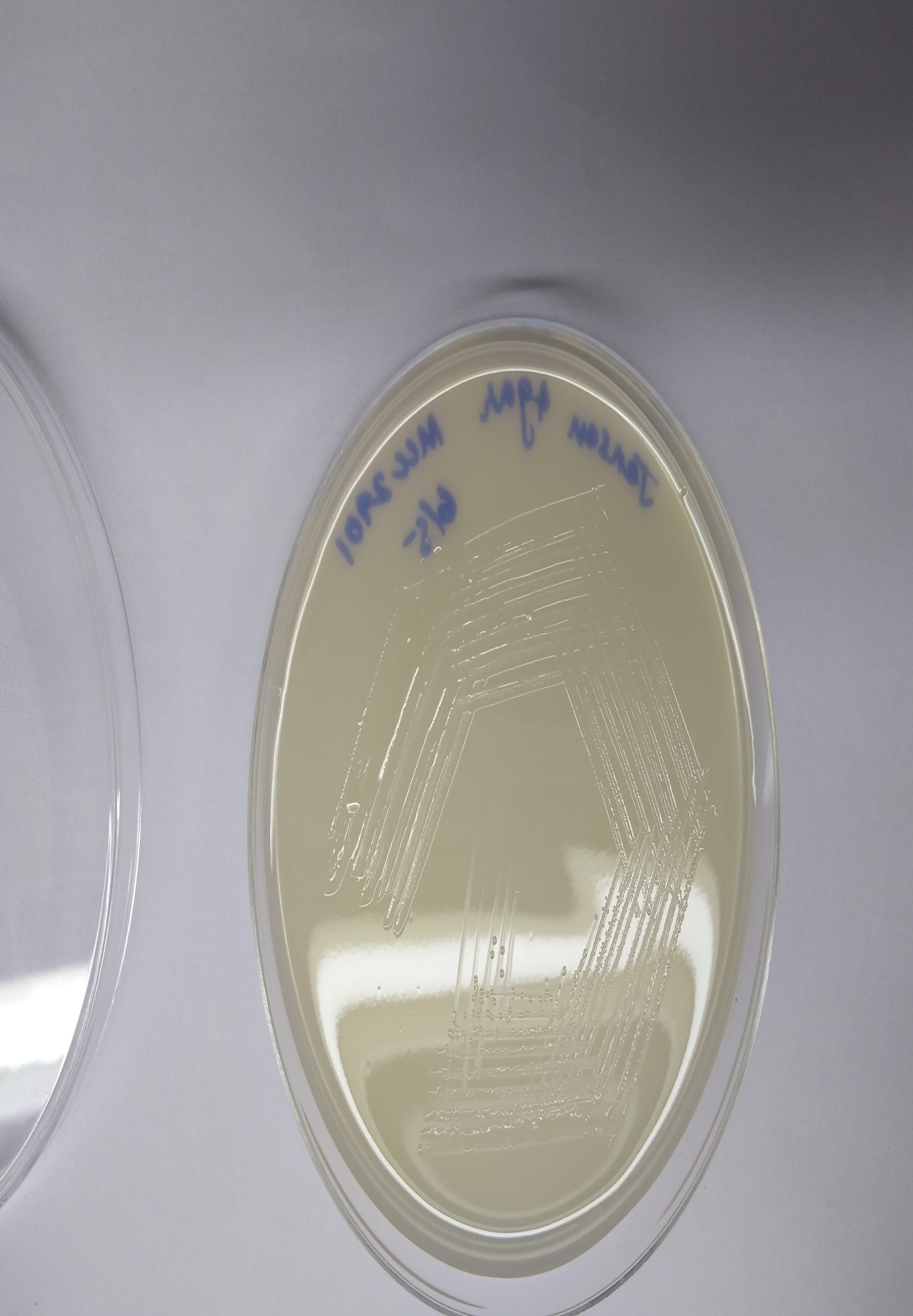

(c)

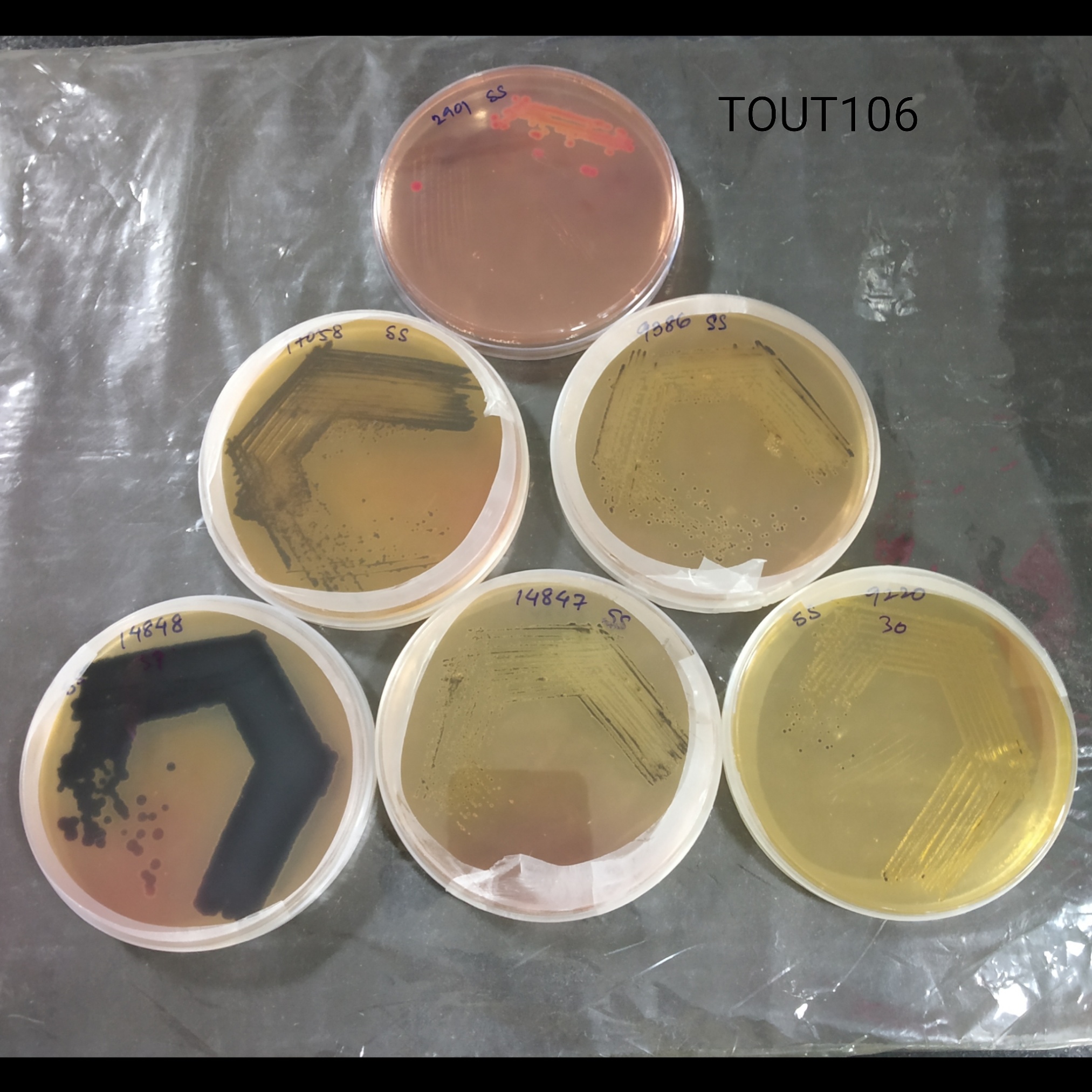

(d)

Fig. S4. Colonies of strain TOUT106^T^ after 24 hrs incubation at 28 °C as observed on (a)MacConkey’s Agar medium; (b) Jensen’s Agar medium; (c) Sheep Blood Agar Medium; and (d) *Salmonella-Shigella* Agar Medium

Table S2 Phenotypic tests of strain TOUT106^T^ on Biolog GN III plates. 1, positive; 0, negative; 0.5, variable.

| Carbon source | TOUT106^T^ |
| --- | --- |
| Dextrin | 0.5 |
| D-maltose | 1 |
| D-trehalose | 1 |
| D-cellobiose | 1 |
| D-gentiobiose | 1 |
| Sucrose | 1 |
| D-turanose | 0 |
| Stachyose | 0 |
| D-raffinose | 1 |
| α-D-lactose | 1 |
| D-melibiose | 1 |
| β-methyl-D-glucoside | 1 |
| D-salicin | 1 |
| N-acetyl-D-glucosamine | 1 |
| N- acetyl-β-D-mannosamine | 0 |
| N-acetyl-D-galactosamine | 1 |
| N-acetyl neuraminic acid | 0 |
| α-D-glucose | 1 |
| D-mannose | 1 |
| D-fructose | 1 |
| D-galactose | 1 |
| 3-methyl glucose | 0 |
| D-fucose | 0 |
| L-fucose | 1 |
| L-rhamnose | 1 |
| Inosine | 1 |
| D-sorbitol | 1 |
| D-mannitol | 1 |
| D-arabitol | 1 |
| Myo-inositol | 1 |
| Glycerol | 1 |
| D-glucose-6-PO_4_ | 1 |
| D-fructose-6-PO_4_ | 1 |
| D-aspartic acid | 0 |
| D-serine | 1 |
| Gelatin | 0 |
| Glycyl-L-proline | 0.5 |
| L-alanine | 0 |
| L-arginine | 0 |
| L-aspartic acid | 1 |
| L-glutamic acid | 0.5 |
| L-histidine | 0 |
| L-pyroglutamic acid | 0 |
| L-serine | 0.5 |
| Pectin | 1 |
| D-galacturonic acid | 1 |
| L-galactonic acid lactone | 1 |
| D-gluconic acid | 1 |
| D-glucuronic acid | 1 |
| Glucuronamide | 0 |
| Mucic acid | 1 |
| Quinic acid | 0 |
| D-saccharic acid | 1 |
| P-hydroxy-phenylacetic acid | 0 |
| Methyl pyruvate | 1 |
| D-lactic acid methyl easter | 0 |
| L-lactic acid | 1 |
| Citric acid | 1 |
| α-keto-glutaric acid | 1 |
| D-malic acid | 1 |
| L-malic acid | 1 |
| Bromo-succinic acid | 1 |
| Tween 40 | 0 |
| G-amino-butryric acid | 0 |
| α-hydroxy-butyric acid | 0 |
| α-hydroxy-D,L Butyric Acid | 0 |
| α-keto-butyric acid | 0 |
| Acetoacetic acid | 0 |
| Propionic acid | 0 |
| Acetic acid | 0 |
| Formic acid | 0 |
| Chemical sensitivity |  |
| pH 6 | 1 |
| pH 5 | 0.5 |
| NaCl 1% | 1 |
| NaCl 4% | 1 |
| NaCl 8% | 0.5 |
| Sodium lactate 1% | 1 |
| Fusidic acid | 0 |
| d-serine | 0 |
| Troleandomycin | 0.5 |
| Rifamycin sv | 1 |
| Minocycline | 0 |
| Lincomycin | 0 |
| Guanidine HCl | 0.5 |
| Niaproof 4 | 1 |
| Vancomycin | 1 |
| Tetrazolium violet | 1 |
| Tetrazolium blue | 1 |
| Nalidixic acid | 0 |
| Lithium chloride | 0.5 |
| Potassium tellurite | 0 |
| Aztreonam | 0 |
| Sodium butyrate | 0.5 |
| Sodium bromate | 0 |

Table S3 Antibiotic susceptibility of strain TOUT106 to the following tested antibiotics by the disk diffusion method as per CLSI guidelines. R: Resistant, S: Susceptible, #Zone diameter interpreted on the basis of standard zones measured for *Escherichia coli* ATCC 25922

| **Antibiotic** | **Concentration (mcg)** | **Zone diameter (mm)** | **Interpretation** |
| --- | --- | --- | --- |
| Amoxicillin | 10 | 0 | R |
| Augmentin | 30 | 0 | R |
| Cefpodoxime | 10 | 11 | R |
| Cephtriaxone | 10 | 33 | S |
| Amikacin | 30 | 20 | S |
| Streptomycin | 10 | 21 | S |
| Gentamicin | 10 | 22 | S |
| Ciprofloxacin | 5 | 31 | S |
| Levofloxacin | 5 | 27 | S |
| Norfloxacin | 10 | 29 | S |
| Chloramphenicol | 30 | 28 | S |
| Tetracycline | 30 | 17 | S |
| Rifampicin^#^ | 5 | 14 | S |
| Cefixime^#^ | 5 | 31 | S |
| Cotrimazole^#^ | 25 | 28 | S |
| Colistin^#^ | 10 | 14 | S |
| Furazolidone^#^ | 50 | 13 | R |
| Netillin^#^ | 30 | 20 | R |

Table S4 MIC of antibiotics for strain TOUT106^T^. R: Resistant, S: Susceptible

| Antibiotic | Minimum Inhibitory Concentration (mcg /ml) | Interpretation |
| --- | --- | --- |
| Colistin | 64 | R |
| Ampicillin | 2 | S |
| Cefotaxime | <1 | S |
| Ciprofloxacin | <1 | S |

Table S5 Enzyme activity of strain TOUT106^T^ using API-Zym test strip. +, positive; -, negative.

| **Enzyme assayed for** | TOUT106^T^ |
| --- | --- |
| Control | - |
| Alkaline phosphatase | **+** |
| Esterase | - |
| Esterase lipase | - |
| Lipase | - |
| Leucine acrylamidase | **+** |
| Valine acrylamidase | - |
| Cysteine acrylamidase | - |
| Trypsin | - |
| a-chymotrypsin | - |
| Acid phosphatase | **+** |
| Naphthol AS-BI-phosphohydrolyase | **+** |
| á-galactosidase | - |
| ß-galactosidase | **+** |
| ß-glucournidase | - |
| á-glucosidase | - |
| glucosidase | - |
| N-acetyl-ß-glucosaminidase | - |
| á-mannosidase | - |
| á-fucosidase | - |
